# Optimization of genomewide CRISPR screens using AsCas12a and multi-guide arrays

**DOI:** 10.1101/2022.10.31.514561

**Authors:** Sakina Petiwala, Apexa Modi, Tifani Anton, Erin Murphy, Sabah Kadri, Hengcheng Hu, Charles Lu, Michael J Flister, Daniel Verduzco

## Abstract

Genomewide loss-of-function (LOF) screening using CRISPR (clustered regularly interspaced short palindromic repeats) has facilitated the discovery of novel gene functions across diverse physiological and pathophysiological systems. A challenge with conventional genomewide CRISPR/Cas9 libraries is the unwieldy size (60,000 to 120,000 constructs), which is resource intensive and prohibitive in some experimental contexts. One solution to streamlining CRISPR screening is by multiplexing two or more guides per gene on a single construct, which enables functional redundancy to compensate for suboptimal gene knockout by individual guides. In this regard, AsCas12a (Cpf1) and its derivatives, e.g., enhanced AsCas12a (enAsCas12a), have enabled multiplexed guide arrays to be specifically and efficiently processed for genome editing. Prior studies have established that multiplexed CRISPR/Cas12a libraries perform comparably to the larger equivalent CRISPR/Cas9 libraries, yet the most efficient CRISPR/Cas12a library design remains unresolved. Here, we demonstrate that CRISPR/Cas12a genomewide LOF screening performed optimally with three guides arrayed per gene construct and could be adapted to robotic cell culture without noticeable differences in screen performance. Thus, the conclusions from this study provide novel insight to streamlining genomewide LOF screening using CRISPR/Cas12a and robotic cell culture.

## INTRODUCTION

CRISPR/Cas9 screening has revolutionized the discovery of novel gene functions across various cellular contexts in normal and diseased states (*1, 2*). Genomewide loss-of-function (LOF) CRISPR/Cas9 libraries have historically included 4-to-6 guides per gene, resulting in libraries ranging from 60,000-to-120,000 constructs and requiring hundreds-of-millions cells per screen (*2–13*). The unwieldy size and resource intensive constraints of genomewide libraries has prompted the miniaturization of a human genomewide CRISPR/Cas9 library with only two optimal guides per gene that performed similarly to larger conventional genomewide libraries (*14*). Notably, this minimal genomewide library still requires ~40,000 constructs and the *in silico* optimization of most guides has not been empirically tested, since the majority of genes are not observable dependencies in >1,000 genomewide LOF studies to date (*14, 15*). Moreover, the ability to overcome variance due to guide-level noise is further compromised by restricting gene-level representation to only two independent guide constructs per gene.

Another approach to reducing CRISPR library size is by multiplexing two or more guides per gene on a single construct, which enables functional redundancy to compensate for suboptimal gene knockout by individual guides. Multiplexing CRISPR/Cas9 guides on a single construct is challenging in that it requires multistep cloning for library construction (16) and is prone to guide uncoupling through recombination (*17, 18*). In contrast, the alternative Cas12a (Cpf1) is distinct in its ability to specifically and efficiently process multiplexed guide arrays for genome editing (*19–21*). Because of this feature, multiple Cas12a guides in an array are cloned in front of a single promoter and expressed as a single transcript. In addition to being highly specific, an enhanced AsCas12a protein (i.e., enAsCas12a) has also been engineered for increased efficiency and greater flexibility to use additional PAM sites (*22, 23*). Finally, Cas12a has recently been adapted for pooled genetic screens with multiplexed guide arrays that perform comparably to traditional CRISPR/Cas9 genomewide LOF libraries (*24, 25*), yet there remain several unaddressed questions regarding the optimal library design.

Despite the advances in CRISPR/Cas12a multiplexed screening, the optimal design remains unclear for genomewide LOF libraries with guide arrays of two or three guides-per-gene in the context of AsCas12a or enAsCas12a. To address this question, a novel 3-guide genomewide LOF library (Cas12-WGx) that is compatible with AsCas12a and enAsCas12a was compared head-to-head with commonly used genomewide LOF libraries for Cas9 (Brunello) and enAsCas12a (Humagne) (*25*). Additionally, the compatibility of the 3-guide library with AsCas12a and enAsCas12a enabled a genomewide head-to-head comparison between the two enzymes for the first time. Finally, to assess the ability to automate CRISPR/Cas12a screening using robotics, the performance of the 3-guide library with AsCas12a was compared between manual screening and robotic screening using the Compact SelecT system. Collectively, these data demonstrated that robotic CRISPR/Cas12 screening performed comparably to manual screening, and the 3-guide multiplexed library with enAsCas12a performed closest to the Brunello CRISPR/Cas9 library and better than all other conditions.

## RESULTS & DISCUSSION

Multiple reports have compared genomewide LOF library designs for Cas9 and Cas12a (*24, 25*), yet no prior studies have directly compared the performance of CRISPR/Cas12 libraries with arrays of two or three guides-per-gene and across both AsCas12a and enAsCas12a enzymes. The Humagne libraries were designed for genomewide compatibility with only the enAsCas12a enzyme and are not backward compatible with wild-type AsCas12a (*25*). To assess the performance across both Cas12 enzymes, a genomewide LOF library targeting 19,701 protein coding genes, with arrays of three guides-per-gene, was designed to be broadly compatible with the AsCas12a and enAsCas12a enzymes. Due to the restrictive canonical PAM sequence of AsCas12a (TTTV), only 18,907 genes (96%) have three TTTV PAM sites, 19,249 genes (98%) have at least two TTTV PAM sites, and 19,493 genes (99%) have at least one TTTV PAM site, whereas 208 genes had no TTTV PAM sites and are therefore not targetable by AsCas12a (**Supplemental Table 1**). For genes with less than three canonical TTTV PAM sites, guides were designed using the alternative PAM sites recognized by enAsCas12a and were added to the library design for a total of three arrayed guides per gene (**Supplemental Table 1**). Thus, all guides were forward compatible with enAsCas12a due to its expanded recognition of TTTV and the three additional PAM sites (*26*). In summary, the Cas12-WGx library enables >99% targeting of protein coding genes within the human genome and with compatibility to use either AsCas12a or enAsCas12a for genomewide LOF screening.

As outlined in **Figure 1**, the primary objective of this study was to benchmark the performance of the genomewide LOF libraries: Cas12-WGx (3-guide arrayed constructs), Humagne (2-guide arrayed constructs), and Brunello (4 independent guide constructs), in the context of AsCas12a, enAsCas12a, or Cas9 enzymes, respectively. Additionally, the Cas12-WGx is compatible with AsCas12a and enCas12a, enabling the comparison of Cas12-WGx performance between the enzymes. Library performance was assessed using NCI-H1299 cells that were engineered to stably express Cas9, AsCas12a, or enAsCas12a enzymes with comparable editing efficiencies, as assessed by an EGFP disruption assay (**Figure 2A**). At 14 days post-transduction of the Cas12-WGx, Humagne, and Brunello libraries in the engineered NCI-H1299 cells, the genomic DNA was extracted, PCR-barcoded, and sequenced at 500X library coverage to deconvolute guide abundances. Principal components analysis (PCA) and Pearson correlation revealed high agreement across library replicates, suggesting that intra-experimental variability would not impact comparisons across libraries and conditions (**Figure 2B** and **Supplemental Figure 1**).

**Figure 1.**
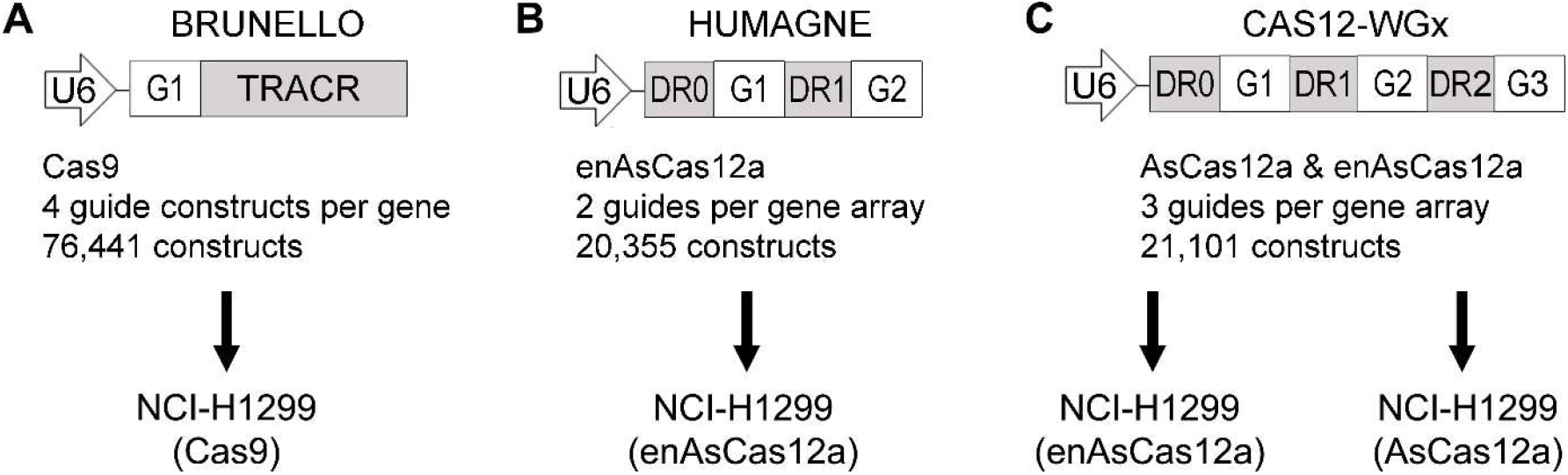
Schematic representation of CRISPR/Cas9 and CRISPR/Cas12 library comparisons in NCI-H1299 cells that were engineered to express spCas9, AsCas12a, or enAsCas12a. (**A**) The Brunello library was designed for spCas9 (PAM = NGG) using Rule Set 2 and consists of 77,441 sgRNAs, with an average of 4 sgRNA constructs per gene and 1,000 non-targeting controls. (**B**) The Humagne D library (guides 2 & 3) was designed for enAsCas12a and consists of 21,820 two-guide arrays per gene, targeting 20,080 protein-coding genes and 1,740 controls. (**C**) The Cas12-WGx library was designed using the Humagne rule sets for AsCas12a and enAsCas12a guides and consists of 20,101 three-guide arrays per gene, targeting 19,701 protein coding genes and 400 controls.

**Figure 2.**
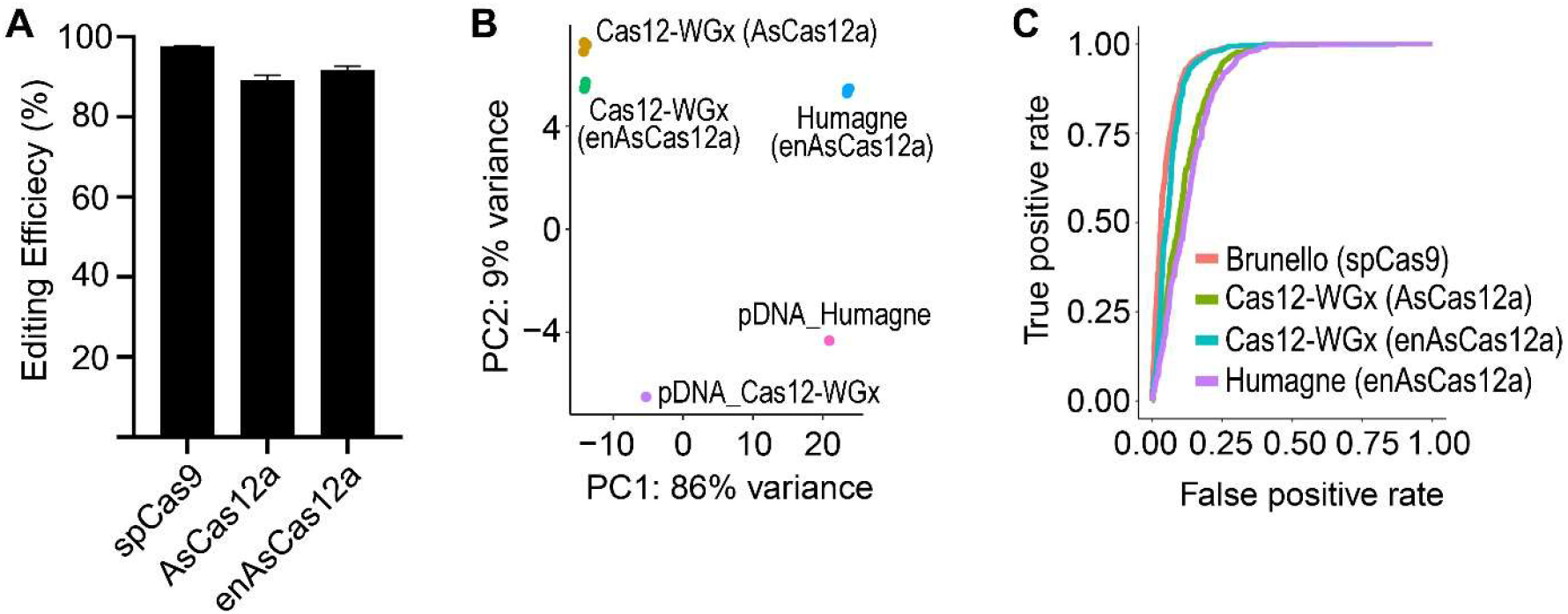
Comparison of library performances for Brunello, Humagne, and Cas12-WGx in 14 day screens using NCI-H1299 cells (n = 3 biological replicates per condition). (**A**) Gene editing activities of spCas9, AsCas12a, and enAsCas12a were assessed in NCI-H1299 cells using an EGFP disruption assay. (**B**) PCA of the Cas12-WGx and Humagne library screens in the context of AsCas12a and enAsCas12a, as indicated on the graph. (**C**) ROC-AUC analysis of Brunello, Cas12-WGx, and Humagne library screens in the context of spCas9, AsCas12a, and enAsCas12a, as indicated on the graph.

Receiver operating characteristic (ROC)–area under the curve (AUC) analysis was used to compare efficacies of Cas12-WGx (asCas12a or enAsCas12a), Humagne (enAsCas12a), and Brunello (spCas9) using genes defined by the DepMap as common essential (true positives) and non-essential (true negatives). Brunello (spCas9) performed well (AUC = 0.95), followed closely by Cas12-WGx with enAsCas12a (AUC = 0.94), whereas the efficacies of Cas12-WGx with AsCas12a (AUC = 0.89) and Humagne with enAsCas12a (AUC = 0.87) were noticeably lower (**Figure 2C**). To further refine the comparison of CRISPR/Cas12a screens, the ROC-AUC analysis was repeated using the 20% most essential and least essential genes identified by log2 fold changes (log2FC) in guide abundances using the CRISPR/Cas9 (Brunello) data in NCI-H1299, which were highly concordant with the NCI-H1299 data from the Cancer Dependency Map (DepMap) (**Figure 3A**). In the refined ROC-AUC analysis, Cas12-WGx with enAsCas12a continued to perform the best (AUC = 0.92), while the lower efficacies remained for Cas12-WGx with AsCas12a (AUC = 0.84) and Humagne with enAsCas12a (AUC = 0.82) (**Figure 3B**). Finally, a precision-recall analysis of the three CRISPR/Cas12a screens using the CRISPR/Cas9 (Brunello)-defined list of essential and non-essential genes in NCI-H1299, also demonstrated that the Cas12-WGx library with enAsCas12a was the most sensitive in measuring gene essentiality (**Figure 3C**). Thus, although all libraries performed within an acceptable range for genomewide LOF screening, these data preliminarily suggested that a 3-guide arrayed library design with enAsCas12a has the highest screening efficacy.

**Figure 3.**
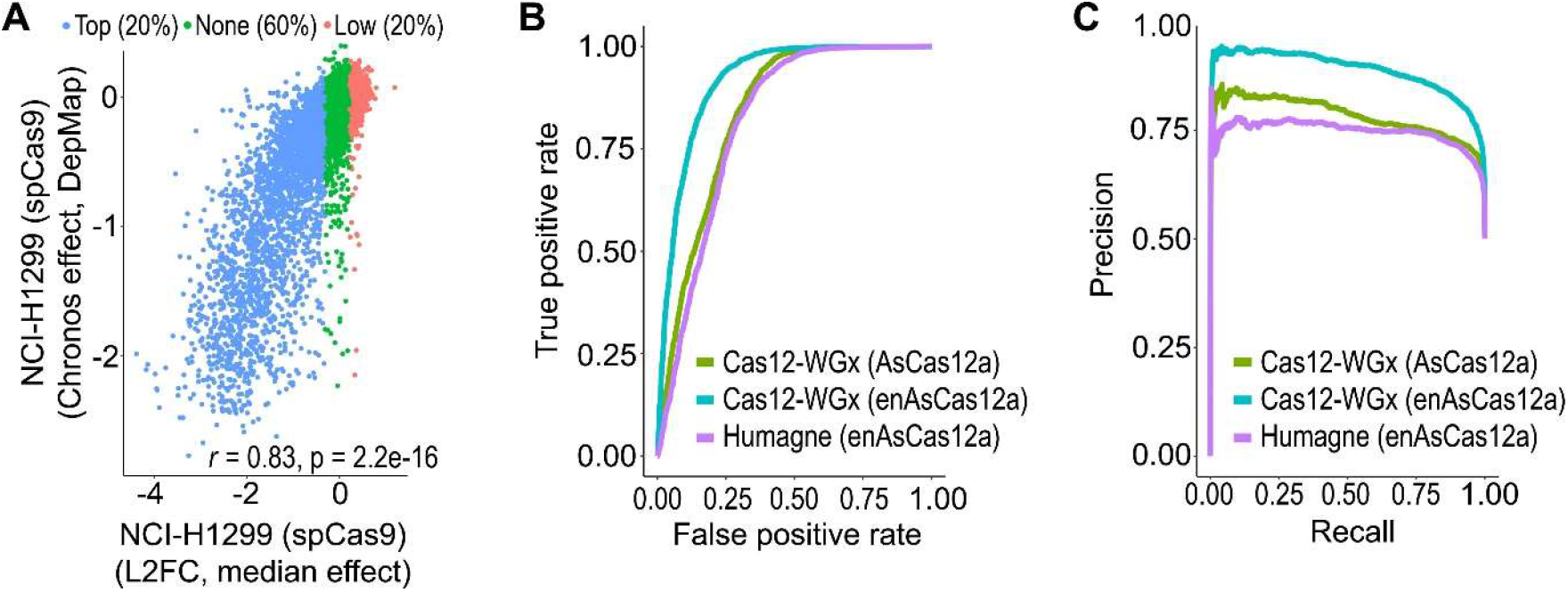
Refined analysis of performance of the Cas12-WGx and Humagne libraries using the essential and non-essential genes identified from Brunello screen in NCI-H1299. (**A**) Comparison of CRISPR/Cas9 (Brunello) data in NCI-H1299 in this study with the NCI-H1299 data from the Cancer Dependency Map (DepMap), revealed high concordance across screens. The top 20% most essential genes (blue) and least essential genes (red) were used for ROC-AUC and precision recall analyses. (**B**) ROC-AUC analysis of Cas12-WGx and Humagne library screens in the context of AsCas12a and enAsCas12a, as indicated on the graph. (**C**) Precision-recall analysis of Cas12-WGx and Humagne library screens in the context of AsCas12a and enAsCas12a, as indicated on the graph.

The observation that enAsCas12a performed better with Cas12-WGx (AUC = 0.94) than Humagne (AUC = 0.87) in NCI-H1299, prompted a head-to-head comparison of the guide arrays shared by both libraries. Since Humagne only has two guides per array, the increased performance of Cas12-WGx could be attributed to the addition of the third guide in the Cas12-WGx library. Comparison of the Cas12-WGx and Humagne libraries revealed 433 guide arrays with two overlapping guides, of which 78 guide pairs targeted essential genes and 78 guide pairs targeted non-essential genes, as defined using the CRISPR/Cas9 (Brunello) data from NCI-H1299 (**Figure 4A**). Using the 156 guide pairs for ROC-AUC analysis revealed that Cas12-WGx performance (AUC = 0.91) was slightly higher than Humagne performance (AUC = 0.89), suggesting that the addition of a third guide to a multi-guide array makes a small and potentially meaningful gain in library performance (**Figure 4B**). Likewise, when considering the arrays with one overlapping guide between Cas12-WGx and Humagne (n = 4,864) (**Figure 4C**), the ROC-AUC analysis revealed that Cas12-WGx (AUC = 0.92) continued to slightly outperform Humagne (AUC = 0.88) (**Figure 4D**), albeit the overall performance of both libraries remained above the level acceptable for genomewide LOF screening.

**Figure 4.**
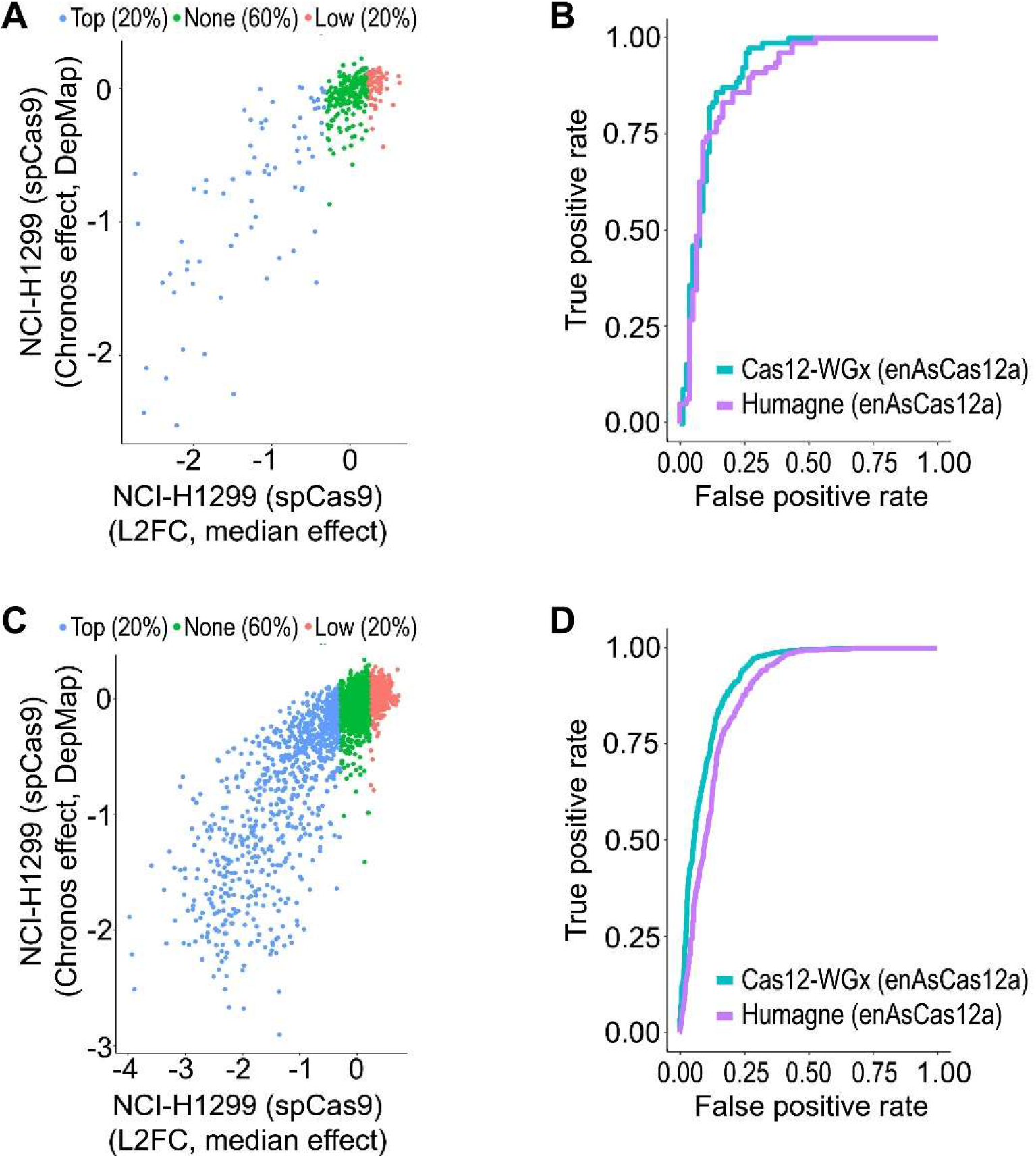
Comparison of Cas12-WGx and Humane library performance using only constructs with one or two guides in common between libraries, and in the context of enAsCas12a only. (**A**) The Cas12-WGx and Humagne libraries share 433 guide arrays with two overlapping guides, from which guide pairs targeting the 20% most essential genes (n = 78) and 20% least essential genes (n = 78) were identified using the CRISPR/Cas9 (Brunello and DepMap) data from NCI-H1299. (**B**) ROC-AUC analysis of the 156 guide pairs shared by Cas12-WGx and Humagne. (**C**) The Cas12-WGx and Humagne libraries share 4,864 guide arrays with at least one overlapping guide, from which guide pairs targeting the 20% most essential genes (n = 965) and 20% least essential genes (n = 1,019) were identified using the CRISPR/Cas9 (Brunello and DepMap) data from NCI-H1299. (**D**) ROC-AUC analysis of the 1,9834 guide pairs shared by Cas12-WGx and Humagne.

A final objective of this study was to test whether genomewide LOF screening with CRISPR/Cas12a could be streamlined further using robotic tissue culture. To answer this question, the Cas12-WGx library was compared in parallel screens using manual cell culture or the Compact SelecT robotic cell culture system from Sartorius (**Figure 5A**). Briefly, the same passage of NCI-H1299 cells expressing AsCas12a were transduced with the Cas12-WGx library, followed by puromycin selection before splitting to replicate flasks that were then cultured manually or robotically for 14 days. At both timepoints, guide abundances were highly congruent between manually and robotically cultured cells, as measured by Pearson correlation (**Figure 5B**) and PCA (**Figure 5C**). Finally, the L2FC in guide abundance were compared between manual and robotic screens, revealing a high concordance between both screening methods (*r* = 0.924) and demonstrating that robotic tissue culture performed comparably to traditional manual screening (**Figure 5D**).

**Figure 5.**
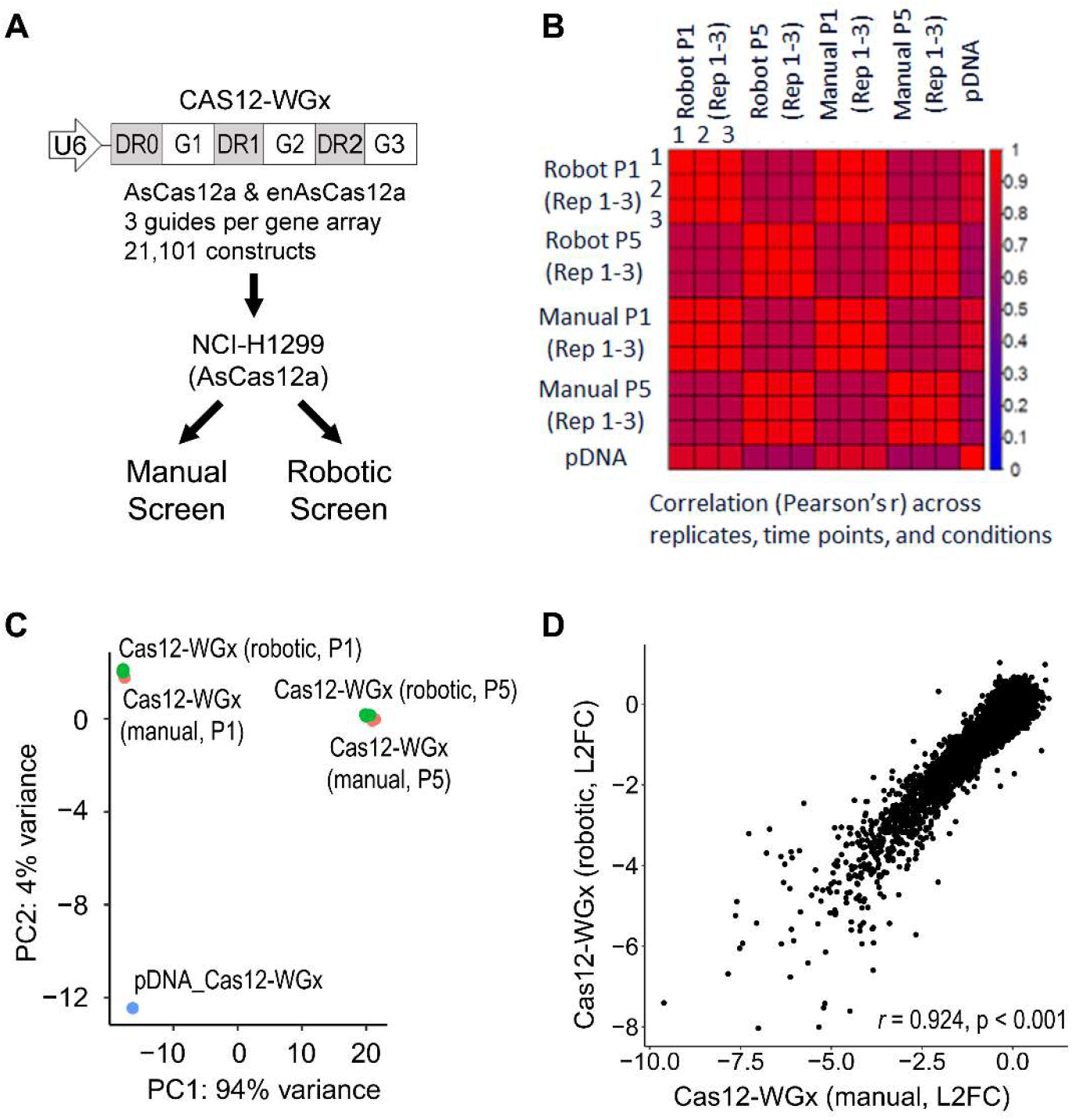
Comparison of Cas12-WGx library performance in manual or robotic 14-day screens using NCI-H1299 cells (n = 3 biological replicates per condition). (**A**) Schematic representation of Cas12a Cas12-WGx screen designs using manual cell culture or the SelecT robotic cell culture system. (**B**) Pearson’s correlation of Cas12-WGx library representation at passages one (P1) and five (P5) using either manual or robotic cell culture. (**C**) PCA of Cas12-WGx library representation at P1 and P5. using either manual or robotic cell culture. (**D**) Pearson’s correlation of the log2 fold chances (L2FC) of guide abundances at the gene level between late and early passages of the manual and robotic cell culture screens.

In summary, this study was the first to demonstrate that CRISPR/Cas12a genomewide LOF screening performed optimally with three guides arrayed per gene construct and could be adapted to robotic cell culture without noticeable differences in screen performance. Moreover, Cas12-WGx with AsCas12a performed comparably to Humagne with enCas12a, suggesting that one additional guide per array compensates for the lower enzymatic activity of AsCas12a (*26*), which would potentially have the added benefit of higher specificity relative to enAsCas12a (*25*). Importantly, we caution against overinterpreting the findings of this study to conclude that the Brunello and Humagne libraries are subpar, as both libraries were highly efficacious and likely well-powered to detect changes with multiple independent guide constructs per gene (*12, 25*). Rather, these data collectively suggest that the most efficient approach to genomewide LOF screening is using a minimal CRISPR/Cas12a library with three guides arrayed per a single gene construct (e.g., Cas12-WGx) and robotic cell culture. Finally, as the design and synthesis of CRISPR/Cas12a libraries continue to evolve, future studies are warranted to assess whether meaningful gains in library performance will be made by multiplexing additional guides per gene array.

## MATERIALS AND METHODS

### Cell Culture

NCI-H1299 cells were cultured in RPMI Medium 1640 (ThermoFisher Scientific #22400071) with 10% fetal bovine serum (FBS, ThermoFisher Scientific # 10082147) and 100U/ml penicillin/streptomycin (ThermoFisher Scientific #15140122). As described previously (*25*), NCI-H1299 cells were engineered to stably express spCas9, AsCas12a, or enAsCas12a enzymes by lentiviral transduction with pXPR_111, pRDA_112, and pRDA_174, respectively. Following transductions, cells were cultured in growth media with blasticidin (10ug/ml for Cas9 cells, 20ug/ml for AsCas12a and 5ug/ml for enAsCas12a cells) and maintained in exponential phase growth by routine passaging in a 37 °C humidity-controlled incubator with 5.0% carbon dioxide. Routine testing for Mycoplasma was performed using a luminescence-based mycoplasma detection assay (Lonza #LT07-705).

### EGFP disruption assay

Parental and Cas9/12a expressing cells were infected with pXPR_011v2/pRDA_221 lentivirus respectively that carries EGFP reporter sequence and a sgRNA targeting EGFP by spinfection in 12 well plates at 1000xg for 2hrs in presence of 10ug/ml polybrene at an MOI of 0.3. After an overnight incubation at 37°C, cells were selected for 3 days in 2ug/ml puromycin. After selection, cells were cultured in presence of puromycin for another 3-4 days and then analyzed for EGFP expression using flow cytometry.

### Library designs

The designs of the Brunello and Humagne libraries have been described in detail elsewhere (*12, 25*). Briefly, the Brunello library was designed for spCas9 (PAM = NGG) using Rule Set 2 and consists of 77,441 sgRNAs, with an average of 4 sgRNA constructs per gene and 1,000 non-targeting controls (*9, 12*). Likewise, the Humagne library design used multiple machine learning approaches to identify optimal enAsCas12a guides by PAM tiling to calculate on-target activity and mismatch analysis to calculate off-target biases using the cutting frequency determination (CFD) score (*25*). Thus, the Humagne library sets (A-D) were designed for enAsCas12a, and each consist of 21,820 two-guide arrays per gene, targeting 20,080 proteincoding genes and 1,740 controls, which were rank-ordered with 1:1 weightings of on-target / off-target scores. For this study, the Humagne Set D (guides 2 & 3) were chosen as an average representation of the performance of the Humagne libraries.

The Cas12-WGx library was designed using the Humagne rule sets for predicting on-target and off-target scores of AsCas12a and enAsCas12a guides (*25*), with the following distinctions: (1) to ensure broad AsCas12a compatibility, preference was given first to guides with TTTV PAM sites, (2) subsequently, enAsCas12a guides (PAMs: TTCC, ATTA, and GTTA) were substituted in the absence of AsCas12a sites passing QC criteria, and (3) guide picking was rank-ordered with 2:1 weightings of on-target / off-target scores. The final Cas12-WGx library consisted of 20,101 three-guide arrays per gene, targeting 19,701 protein coding genes and 400 controls. Guide arrays were designed to express as contiguous transcripts with 20nt direct repeat (DR) sequences and 23nt guide sequences interspersed as follows: DR0-sgRNA1-DR1-sgRNA2-DR2-sgRNA3. The library oligo pools were synthesized by TWIST Biosciences and cloned into pRDA_052 by the Genetic Perturbation Platform (GPP) team at the Broad Institute, as described previously (*25*).

### Genomewide pooled CRISPR screens

As described previously (*25*), NCI-H1299 that were engineered to express spCas9, AsCas12a, or enAsCas12a were transduced with the Brunello, Cas12-WGx or Humagne sgRNA library, respectively. Briefly, lentiviral guide libraries were titrated to ensure low multiplicity of infection (MOI < 0.5) in NCI-H1299 cells at 500X library coverage using 10ug/ml of polybrene. At 24h post-transduction, fresh media was replaced with 2ug/ml puromycin and 10ug/ml of blasticidin and cells were cultured for 14 days by passaging every 3-4 days to maintain exponential phase growth. For each library, cells were transduced in triplicate and cultured independently at a representation of 500cells/guide. At day 14, cells were pelleted by centrifugation and frozen at -80°C for genomic DNA extraction.

### Genomic DNA extraction, PCR barcoding, and NextGen sequencing

Genomic DNA (gDNA) extraction, PCR barcoding, and NextGen sequencing (NGS) were performed, as described previously (*25*). Briefly, gDNA was extracted using a Nucleospin Blood Maxi kit (Takara, Cat. #740950) (for CRISPR/Cas9 screening) and Nucleospin Blood Midi kit (Takara Cat. #740954) (for CRISPR/Cas12a screening), per manufacturer guidelines. The extracted gDNA were PCR barcoded using the following master mix (100μL reactions): 10 μg of gDNA, 10μL of 10x Titanium *Taq* PCR Buffer, 1.5μL Titanium *Taq* DNA Polymerase (Clontech Takara, Cat# 639242), 8 μL dNTPs (Clontech Takara Cat# 4030), 5μL DMSO (Sigma Aldrich Cat# D9170-5VL), 0.5μL P5 primer mix (100μM), 10μL of P7 primer (5μM), and molecular grade H2O (q.s. to 100μL). PCR cycling was performed as follows: an initial denaturation step at 95 °C (5min); followed by 28 cycles of 95 °C (30s), 53 °C (30s), and 72 °C (20s); and a final extension at 72 °C (10min). PCR primers were synthesized at Integrated DNA Technologies and the sequences for Argon P5 and Kermit P7 are provided in (**Supplemental Table 2**). To maintain 500X library coverage during PCR barcoding, a total of 24 reactions were performed for each CRISPR/Cas9 screen replicate and 6 reactions for all CRISPR/Cas12a screen replicates. The PCR barcoded library amplicons were purified using Ampure beads per manufacturer’s instructions (Beckman Coulter, A63880). The purified libraries were then sequenced at 500X coverage using 75nt single end reads on a NextSeq 500 (Illumina).

### Screen Analysis

Sequencing data from each CRISPR screen were demultiplexed and reads with guide RNA (sgRNA) sequences were quantified using custom Perl scripts. Given that the position of the sgRNA could vary per read, the primer sequences flanking the sgRNA were considered. Sequences in between the flanking primers were extracted and then compared to sequences in the sgRNA library. Only sequences with no mismatches were used in the calculation of guide-level read counts. Samples with a minimum of 80% reads mapping to a guide RNA were considered for further analysis. Through the edge R Bioconductor package (*27*), reads were normalized between samples using the trimmed mean of M values (TMM) method (*28*). To measure guide heterogeneity across timepoints the gini index for each sample was calculated. Differential analysis of guide level counts was performed using the limma R Bioconductor package (*29*). For the Cas9 screen, gene level effects were considered as the median log fold change of the four guides associated with each gene. In Cas12a screens, all guides per gene are on the same construct and so the guide-level log fold change is synonymous with gene level effect. Statistical tests of the differential analysis were carried out using the Robust Rank Aggregation (RRA) (*30*) method at the gene level.

### Benchmarking CRISPR screen performance

To determine how well screens predict gene essentiality, we trained a logistic regression classifier. We used the logFC and FDR value associated with each gene as input for the classifier and defined the gene essentiality truth labels as genes overlapping DEPMAP common essential (n=1786) and non-essential genes (n=739) (*6, 31*). In a second classifier we defined essential and non-essential gene truth labels as the top and bottom 20% of genes, ranked based on median effect, in our Cas9 screen. We used the train function from the caret package in R (v3.6.1) to train a separate model per screen using 10-fold cross validation. Parameters were set as follows: train(Type~logFC+FDR,data=cas12,trControl=trainControl(method=“cv”,number=10,savePredictions=T,returnData=TRUE,classProbs=T), method = “glm”,family=“binomial”). In this function the dataset is split 10 times, each time into 10 folds with one-fold being assigned as the test set, resulting in 10 test sets. The label probabilities across all 10 folds were saved into a caret model, which was then used directly as input for the Mleval package in R (v3.6.1) which calculates metrics for machine learning model evaluation, including the area under the receiver-operator curve, and precision-recall curves.

## FIGURE LEGENDS

**Supplemental Figure 1.**
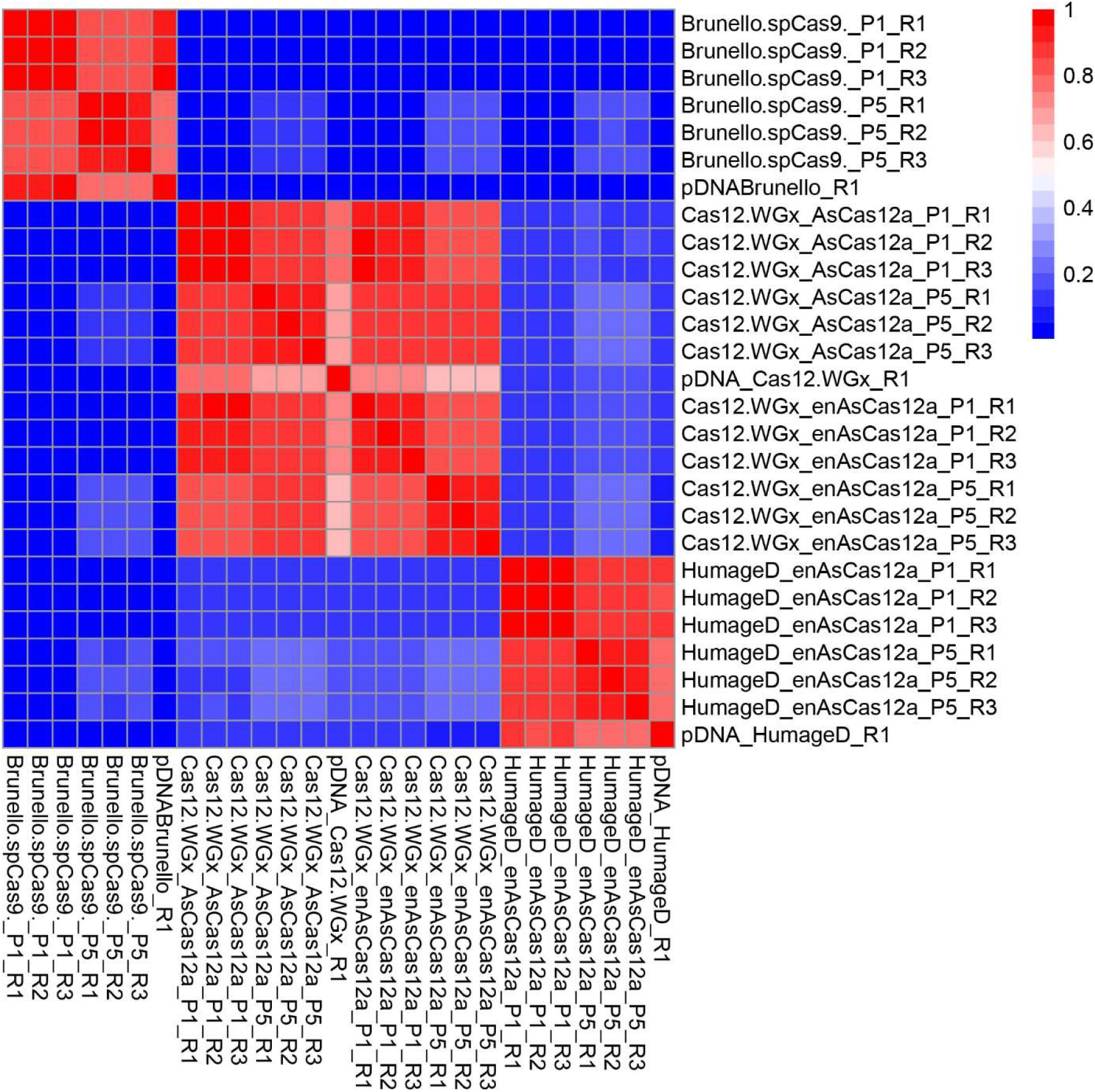
Pearson’s correlation across all libraries and conditions for the manual CRISPR screens. The Pearson’s coefficient (*r*) is indicated by the color key on the right.

